# The proneurogenic and microglial modulatory properties of botulinum toxin in the hippocampus of aging experimental mice

**DOI:** 10.1101/2024.06.14.599127

**Authors:** Jerly Helan Mary Joseph, Mercy Priyadharshini Babu Deva Irakkam, Mahesh Kandasamy

## Abstract

This study explored the neurogenic and microglial modulatory properties of botulinum toxin in the hippocampus of aging experimental mice. Therapeutic botulinum toxin (BoNT) treatment is widely practiced to reduce the excessive discharge of acetylcholine (ACh) in the management of aging and neurological deficits. While the production of new neurons in the adult brain contributes to cognitive functions, age-related diseases with excessive release of ACh and progressive neuroinflammation have been characterized by impaired hippocampal neurogenesis and memory loss. Therefore, we investigated the effect of BoNT on the regulation of hippocampal neurogenesis, focusing on doublecortin (DCX)-positive immature neurons in the hippocampus of aging experimental mice. We also assessed ionized calcium-binding adapter molecule 1 (Iba1)-positive microglia and the expression of cyclooxygenase (COX)-2, a key inflammatory response element, using reverse transcription polymerase chain reaction (RT-PCR). Results revealed a prominent increase in DCX-positive cells in the BoNT-treated animals compared to the control group. Additionally, the reduced number of microglia accompanied by decreased mRNA expression of COX-2 was evident in the BoNT-treated animals. These dual effects suggest that BoNT could be a promising therapeutic agent for mitigating age-related neuroregenerative decline and neuroinflammation responsible for cognitive impairments.

## 1. Introduction

Implementation of a mild dose of botulinum toxin (BoNT) has been a well-established medical practice in the therapeutic management of migraine, neuropathic pain, neuromuscular disorders, and various movement disorders including dystonia, cerebellar ataxia, Parkinson’s disease (PD), and Huntington’s disease (HD) (Nigam and Nigam 2010). We have recently demonstrated that a single shot of intramuscular injection of BoNT reduces anxiety, and improves learning and memory in experimental aging mice (Yesudhas et al. 2020, 2021). The continual generation of new neurons in the hippocampus of the adult brain has been attributed to the cellular basis of mental ability, mood, and cognitive functions in healthy situations. In contrast, aging, chronic stress, anxiety, and progressive neurodegenerative conditions have been characterized by elevated neuroinflammatory processes and impaired hippocampal neurogenesis accounting for memory loss. Activation of microglia, a cellular entity responsible for innate immune responses in the brain, plays a crucial role in neuroinflammation, which is linked to impaired neurogenesis (Fornari Laurindo et al. 2024). Considering its positive influence on hippocampal plasticity, BoNT treatment could be involved in the regulation of neural regeneration in association with the modulation of microglia in the hippocampus of the adult brain.

Doublecortin (DCX), a microtubule-stabilizing protein essential is essential for neural migration in the developing and adult brains. As DCX is a putative marker of immature neurons and neuroblasts in the adult brain. The number of DCX-positive cells has been considered to represent the number of neurogenic processes in the adult brain (Couillard-Despres et al. 2005). While microglia contribute to the regulation of neuroplasticity and neurogenic processes in the physiological state, their activation during aging and brain diseases contributes to neuropathogenic mechanisms hindering neurogenesis in the hippocampus. The expression of ionized calcium-binding adapter-1 (Iba-1) has been ascertained as a valid marker of microglia in the brain. While overexpression of cyclooxygenase (COX)-2, a molecular indicator of inflammation is known to activate microglia, activated microglia serve as a potential source for the elevated levels of COX-2 thereby, both can synergistically exert deleterious effects on hippocampal neurogenesis in the brain (Vijitruth et al. 2006; Nagano et al. 2021). Taken together, the observed BoNT-mediated effects on cognitive improvement and anxiolytic properties might be associated with the alteration in immature neuronal and microglia populations. Therefore, this study examined the influence of therapeutic BoNT on the regulation of hippocampal neurogenesis at the immunohistochemical levels of DCX and Iba-1-expressing cells. Additionally, the expression of COX-2 was assessed using reverse transcription polymerase chain reaction (RT-PCR) in the hippocampus of aging experimental brains.

## 2. Materials and methods

### 2.1. Experimental Animals and BoNT Treatment

Male BALB/c mice aged 7-8 months (N=12) were randomly assigned to two groups such as control (N=6) and BoNT (N=6). The BoNT group received 1U of intramuscular injection of therapeutic BoNT (Allergan, Ireland), per KgBW, while the control group received saline. After a month, experimental mice were perfused and the collected brains were processed for cryosections. Similarly, an additional set of mice, control (N=3), and BoNT (N=3) were also sacrificed without perfusion, and their brains were included for molecular biological assessments. The study was approved by the institutional animal ethics committee (IAEC), Bharathidasan University under the direction of the committee for the purpose of control and supervision of experiments on animals (CPCSEA), India (Ref No: BDU/IAEC/P27/2018, August 07, 2018)

### 2.2. Perfusion of experimental animals and brain cryosectioning

The experimental mice were subjected to transcardial perfusion using sterile 0.9% NaCl (Sisco Research Laboratories (SRL) 41721, India) followed by 4% paraformaldehyde (PFA) (Himedia GRM3660-500G, India). The dissected brains were immersed overnight in 4% PFA at 4°C. Subsequently, the brains were transferred to sterile containers with 30% sucrose (SRL 27580, India). After a week, the brains were coated with Poly Freeze medium (Sigma-Aldrich SHH0026, USA) and subjected to 30µm sagittal sections on a sliding microtome (Weswox, India) using dry ice. The serial sections of the brains were collected and stored in tubes filled with a cryoprotectant buffer solution comprising glycerol (SRL 42595, India) and ethylene glycol (Himedia AS052-500ML, India) and maintained at -20 °C for subsequent immunohistochemical analysis.

### 2.3. Immunohistochemical Analyses

For the immunohistochemical analysis, every twelfth section, spaced 360µm apart, from a hemisphere was selected and placed in a twelve-well plate (Tarson, India) and washed with 1x Tris-buffered saline (TBS) for 10 minutes x 3 times using an orbital shaker (Tarson, India) at room temperature. Then, the brain sections were subjected to antigen retrieval by incubation with 10 millimolar (mM) Tri-sodium citrate(Thermo Fisher Scientific 27625, USA) in 0.01% Triton X (HiMedia MB031-50ML, India, pH 6.0) in a water bath at 65°C for 2 hours. Following the washing step, the sections were exposed to 3% Bovine serum albumin (BSA) (Himedia GRM3115-100G, India) for 1 hour at room temperature. Immunolabeling of immature neurons and microglia was done by incubating the sections with rabbit anti-DCX antibody (Cell Signalling Technology 4604S, USA; 1:250 dilution) and rabbit anti-Iba-1 antibody (Cell Signaling Technology 17198S, USA, 1:250 dilution) respectively, for 48 hours at 4 °C. Afterward, the sections underwent three consecutive washes with 1x TBS for 10 minutes each and incubated with a goat anti-rabbit secondary antibody conjugated with a fluorescent tag, Dylight 594 (Novus Biologicals NBP1-72732DL594, USA; 1:500 dilution) for 24 hours at 4 °C. On the subsequent day, sections were washed 3 times with 1x TBS for 10 minutes and the sections were arranged on double-frosted microscope slides (Borosil, India), and allowed to dry overnight. The sections on the microscope slides were mounted with coverslips using ProLong™ Glass Antifade Mountant (Thermo Fisher Scientific P36982, USA) solution and dried. Then blind codes were assigned to the specimens and immunolabeled cells were digitally captured using a fluorescence microscope (DM750, Leica Microsystems, Germany). The DCX- and Iba-1-positive cells were quantified in the hippocampal DG with the aid of the ImageJ with cell counter plugin. The total number of DCX- and Iba-1-positive cells was estimated in the hippocampal DG and the average number of cells per hippocampal section was depicted.

### 2.4. RNA Isolation

Using a dissection microscope, the hippocampi were carefully separated from the brains of the second set of experimental animals and homogenized with 200μl of Trizol reagent (Ambion 15596026, USA) using a homogenizer (Merck, Germany). The resulting homogenates were shaken with 200μl of chloroform (Himedia MB109-500ML, India) and incubated at room temperature for 3 minutes. Then the tubes were centrifuged using a cooling centrifuge (REMI, Mumbai, India) at 12,000 rpm for 15 minutes at 4°C. The aqueous phase was transferred to a fresh sterile tube containing 100μl of isopropanol (EMPLURA® 8.18766, Merck, Germany). After thorough mixing, the tubes were incubated for 10 minutes at room temperature and centrifuged at 12,000 rpm for 20 minutes at 4°C. The resulting supernatants were carefully removed and the RNA pellets were washed with 200μl of 75% ethanol. Then the sample tubes were centrifuged at 7,500 rpm for 5 minutes at 4°C and the supernatants were carefully discarded. The RNA pellets were air-dried for 5-10 minutes and dissolved in sterile RNAase-free water. The purity and the concentration of the total RNA were measured based on absorbance readings at 260 nm using spectrophotometry (Eppendorf, Germany) and subjected to RT-PCR.

### 2.5. RT-PCR

For the synthesis of cDNA, 1µg of total RNA was mixed with a total volume of 10µl of cDNA synthesis kit (TaKaRa 6110A, Tokyo) consisting of dNTP mixture, Oligo dT primer, Random 6mers, 5x prime-script buffer, RNAase inhibitor, prime-script RTase, nuclease-free water and reverse-transcribed according to the manufacturer protocol by incubation at 30°C for 10 minutes, 50°C for 60 minutes, and 95°C for 10 minutes using a thermocycler (Veriti, Applied Biosystems, USA). Then 1µl of cDNA was subjected to amplification in a total volume of 10µl reaction mixture containing 2x master mix (Ampliqon A140301, Denmark), Forward and reverse primer, cDNA template, and nuclease-free water (Qiagen 129115, Germany). The PCR conditions for COX-2 are as follows: denaturation at 95°C for 10 minutes, annealing at 58°C for 1 minute for 32 cycles, and extension at 72°C for 10 minutes (Forward primer: 5’ AGAGTCACCACTACGTCA 3’and Reverse primer: 5’ ATCTGGGTCTACCTATGTA 3’). The conditions of PCR for the housekeeping gene, Tubulin are as follows: denaturation at 95°C for 10 minutes, annealing at 60°C for 1 minute for 32 cycles, and extension at 72°C for 10 minutes (Forward primer: 5’ CATCGACAATGAAGCCCTCTA 3’and reverse primer: 5’CTTTAACCTGGGAGCCCTAATG 3’). The PCR products were separated by 2% agarose gel electrophoresis, and stained with ethidium bromide. The gel image was photo-documented using the gel documentation system (Bio-Rad, USA). The digital images of the gel were subjected to the densitometric assessment using ImageJ software. The mRNA expression levels of COX-2 were normalized with the expression of the Tubulin in the control and BoNT-treated groups respectively.

### 2.6. Statistical Analyses

Statistical Analysis was done using GraphPad Prism software. The values were represented as mean ± standard deviation (SD). Student’s *t*-test was used to measure the statistical differences *P*-value < 0.05 is considered to be statistically significant.

## 3. Results

### 3.1. BoNT-treated mice show an increased number of DCX-positive immature neurons in the hippocampal DG

The quantification of the DCX-positive cells in the hippocampal DG region revealed an increased number of immature neurons in the brains of the BoNT-treated experimental mice than that of the control (Control = 55 ± 15 vs BoNT = 89 ± 33) (Fig. 1).

**Fig 1:**
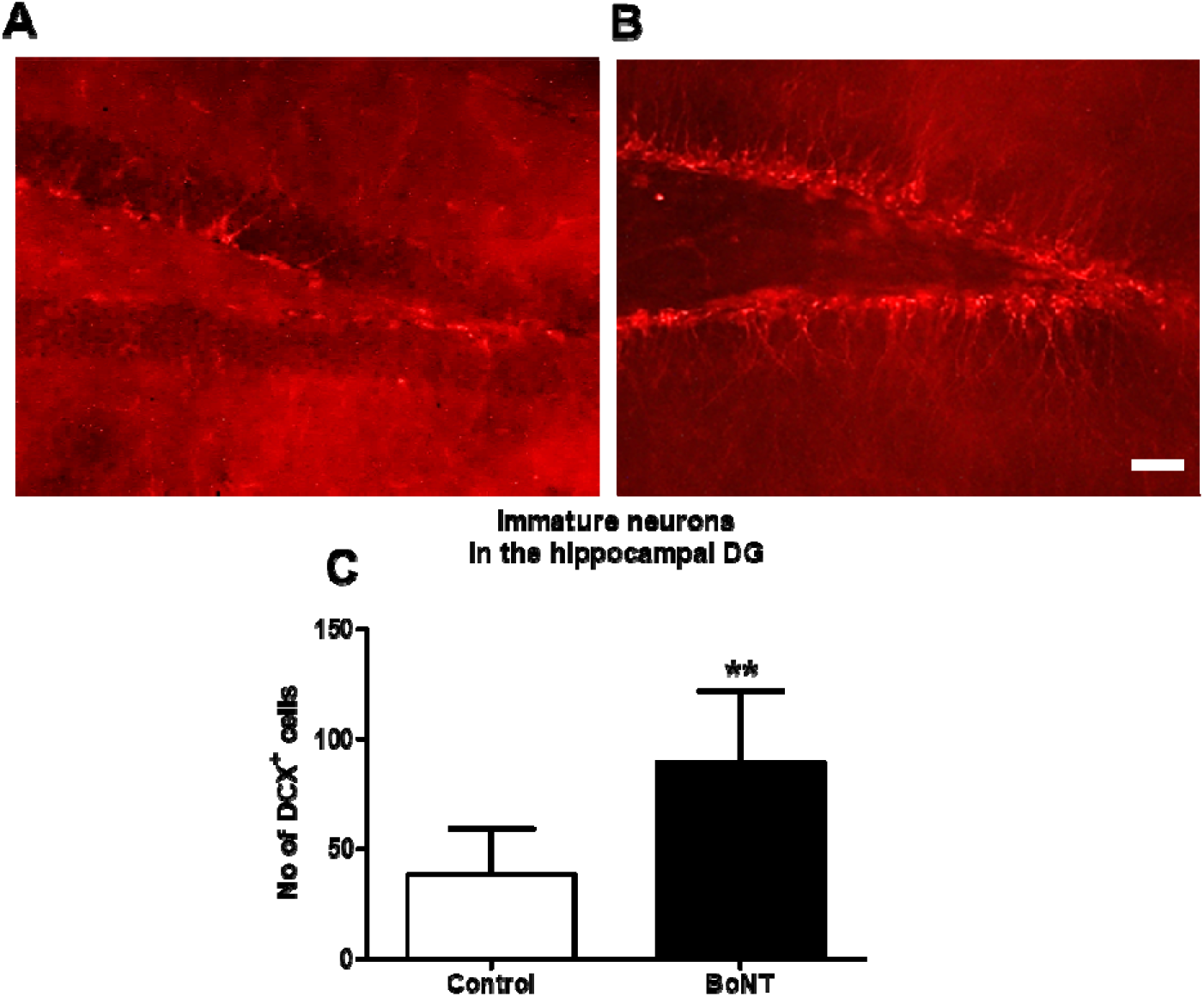
Immunohistochemical analysis of DCX-positive cells. Representative image of DCX immuno-positive cells in hippocampal DG of **A)** control and **B)** BoNT-treated mice. The scale bar = 25µm. **C)** The bar graph represents the number of DCX-positive cells.

### 3.2. BoNT treatment reduced the number of Iba-1-positive microglia in the hippocampal DG of experimental mice

The immunohistochemical quantification of Iba-1 positive cells indicated a significant reduction in the number of microglia in the hippocampal DG of BoNT-treated mice compared to the control group (Control = 52 ± 7 vs BoNT = 34 ± 8) (Fig. 2).

**Fig 2:**
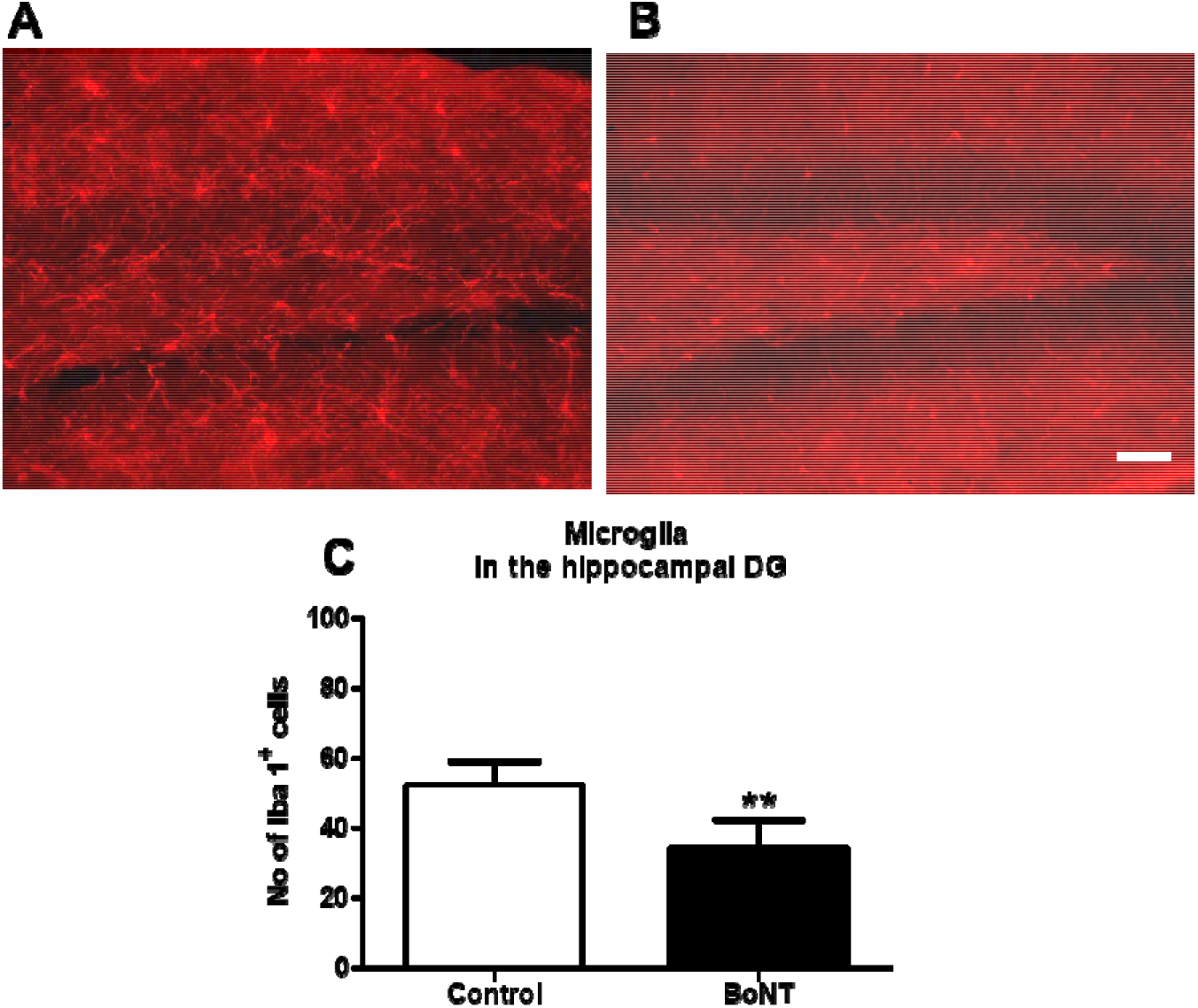
Immunohistochemical analysis of Iba-1 positive cells. Representative image of Iba-1 immunopositive cells in hippocampal DG of **A)** control and **B)** BoNT-treated mice. The scale bar = 25µm. **C)** The bar graph represents the number of Iba-1 positive cells in the hippocampal DG region.

**Fig 3:**
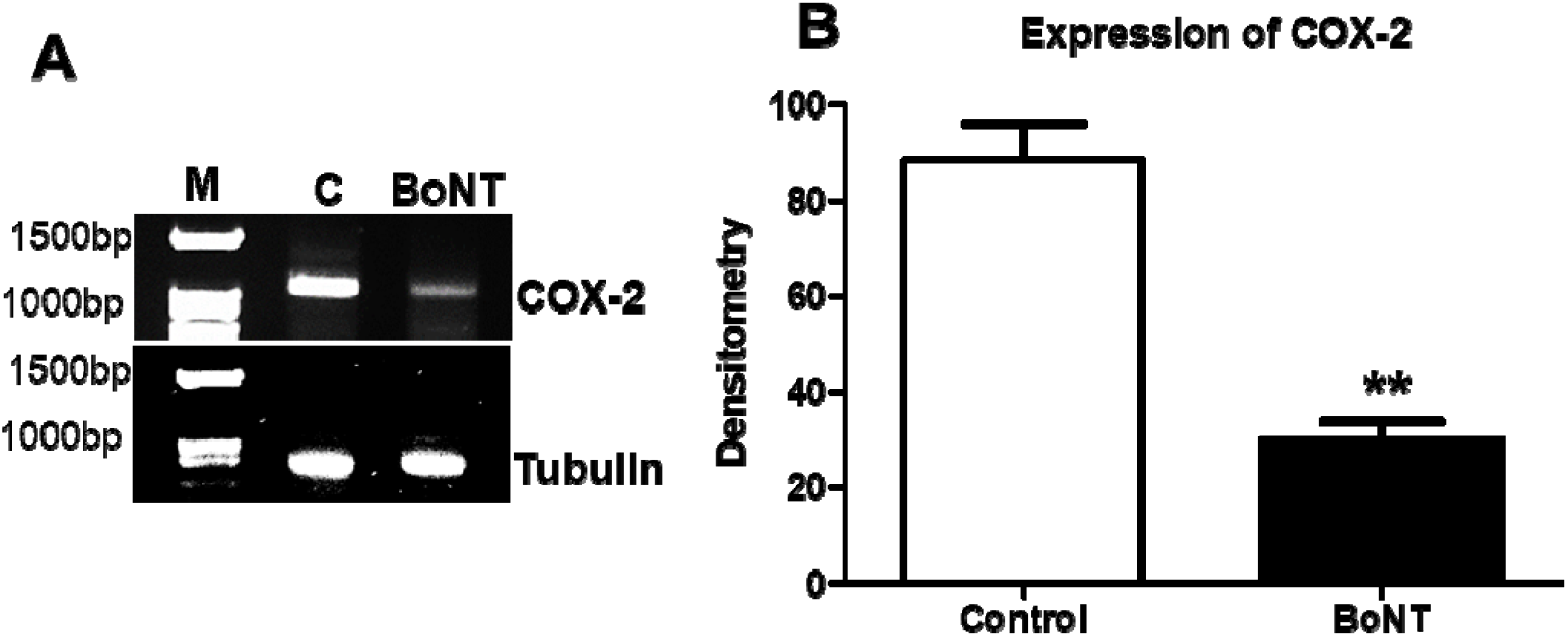
RT-PCR results of Tubulin and COX -2 A) Agarose gel images of the PCR products of COX-2 and Tubulin and B) Bar graph of the densitometric analysis for the same in the hippocampal DG of control and BoNT treated animals.

### 3.3. BoNT treatment reduced the expression of COX-2 in the hippocampal DG of experimental mice

The densitometry assessment of the RT-PCR product indicated a marked reduction in mRNA expression of COX-2 in the hippocampus of the experimental mice BoNT-treated group than the control group (Control = 88 % ± 14 vs BoNT = 30 % ± 7) (Fig.3).

## 4. Discussion

The brain regenerative processes via hippocampal neurogenesis regulation have been suggested to be important for cognitive functions in the physiological states. Aging and neuroinflammation play a detrimental role in suppressing neurogenesis in the hippocampus leading to progressive memory loss, anxiety, and depressive-like symptoms. The aging brain is highly vulnerable to neuroinflammation and oxidative damage, significantly increasing the risk of developing dementia. Thus, pharmacological interventions targeting neuroinflammatory pathways in the brain might aid in the development of potential therapies for neural regeneration.

Recent reports indicate that administering a mild dose of BoNT via intramuscular injection reduces anxiety and enhances learning and memory in experimental aging mice. This effect is considered to result from BoNT-mediated neuroprotection and antioxidant activities within the brain (Yesudhas et al., 2020, 2021). BoNT administration has been reported to suppress the expression of proinflammatory genes in the 1-methyl-4-phenyl-1,2,3,6-tetrahydropyridine (MPTP) and 6-OHDA hydroxydopamine (OHDA-induced experimental models of PD (Ham et al. 2022). Yang Li et al. reported that 3 days of injections of BoNT attenuated depressive-related behaviors in association with reduced activation of microglia in the hippocampus of a reserpine-induced mouse model of PD (Li et al. 2023). An increased level of COX-2 has been linked to the activation of microglia and induction of free radical oxidative damage and impairing the hippocampal neurogenesis brain the present study denotes a significant reduction of microglial cells and COX-2 expression in the hippocampus of the adult brain. A study by Yao-Chi Chuang et al reported that intravesical administration of BoNT inhibited the expression of COX-2 and E-type prostanoid receptor-(EP)-4 which in turn attenuated bladder hyperactivity in cyclophosphamide-induced cystitis in rats (Chuang et al. 2009). Taken together, the present report supports the previous findings that BoNT treatment mediates the reduction of microglia and inhibition of COX-2 expression in various experimental disease conditions. Notably, suppression of neuroinflammation has been proposed to facilitate neurogenesis, synaptic plasticity, and neuroprotection in the brain.

To note, Chiara Panzi et al demonstrated that therapeutic BoNT reduces the release of mutant Tau protein, a causative agent of Alzheimer’s disease, in the primary mouse hippocampal neurons (Panzi et al. 2023). Solabre Valois et al reported that the incubation of hippocampal neurons with a non-toxic C-terminal region of the receptor-binding domain of heavy chain BoNT promoted the axonal outgrowth of dendritic arborizations via the activation of small Ras-related C3 botulinum toxin substrate (Rac)-1 Rho guanosine triphosphate hydrolases (GTPases) and extracellular signal-regulated kinase (ERK) pathway (Solabre Valois et al. 2021). Hence, beyond modulation of cholinergic functions, BoNT treatment induces non-canonical bioregulatory effects on gene, metabolic, signaling pathways, and cell cycle alterations in the brain. Wang, L reported that following 48 hours of treatment with BoNT, SH-SY5Y cells exhibit induced expression transcripts related to neuronal cell development and metabolic processes (Wang et al. 2020). BoNT has been shown to up-regulate the level of brain-derived neurotrophic factor (BDNF), which is a crucial neurotrophin involved in the differentiation, neuron survival, and synaptic plasticity of newborn neurons in the adult brain (Hwang et al. 2023), Therefore, the increased number of immature neurons observed in the hippocampus of BoNT-treated groups could be attributed to BoNT-mediated neurotrophic support and its anti-neuroinflammatory properties.

Considering the facts, this study supports a hypothesis that the BoNT-mediated increased neurogenic process resulting from its anti-inflammatory, neuronal differentiation, and neuroprotective measures, might collectively underlie its anxiolytic and procognitive effects. However, further studies are required to elucidate the precise molecular mechanisms underlying the neuroregenerative and anti-neuroinflammatory effects of BoNT in the brain.

In summary, the study provides compelling evidence for the potential of therapeutic BoNT to modulate hippocampal neurogenesis and neuroinflammation in the brain, highlighting its evolving role as a promising therapeutic agent for neurological complications with memory loss beyond its traditional neuromuscular applications.

## 5. Conclusion

BoNT treatment facilitates pro-neurogenic and anti-inflammatory effects in the hippocampal DG of experimental aging animals. The enhanced neurogenic process and neuroprotection could serve as an underlying cellular basis for the cognitive improvement and anxiolytic effects of BoNT. Thus, BoNT holds a promise for treating anxiety, neurodegenerative disorders, and dementia.

## Availability of data

Data is provided within the manuscript

## Acknowledgments

This work was supported by a research grant (SERB-EEQ/2016/000639) from the Science and Engineering Research Board (SERB). MK has been supported by the Faculty Recharge Programme, University Grants Commission (UGC-FRP), New Delhi, India. The authors also thank RUSA 2.0, Biological Sciences, Bharathidasan University, for their financial support (TN RUSA: 311/RUSA (2.0)/2018 dt. 2 December 2020).

The authors would like to thank Prof. Anusuyadevi Muthuswamy for their helpful scientific discussions and suggestions on the plan and protocol of the manuscript. Authors acknowledge UGC-SAP and DST-FIST for the infrastructure of the Department of Animal Science, Bharathidasan University.

## Author contribution statement

MK: Conceptualization, project administration, supervision, funding acquisition, data curation, writing-original draft, review, and editing. JHMJ: Writing-original draft, methodology, acquisition of data, validation, formal analysis. MPBDI: methodology, acquisition of data, validation, review

## Ethical approval

This work was revived and approved by the Institutional Animal Ethics Committee (IAEC), Bharathidasan University (Ref No: BDU/IAEC/P27/2018, August 07, 2018), under the regulation of the Committee for the Purpose of Control and Supervision of Experiments on Animals (CPCSEA), India.

